# Early-Life Poultry-Derived Lactobacilli Drive Microbial Succession and Gut Immune Modulation in Broiler Chickens

**DOI:** 10.1101/2025.08.01.668251

**Authors:** Shreeya Sharma, Anna Seekatz, Mohammadali Alizadeh, Hosni Hassan, Alexander Yitabrek, Scott Pratt, Khaled Abdelaziz

**Author notes:** Corresponding author Khaled Abdelaziz Department of Animal and Veterinary Sciences College of Agriculture, Forestry and Life Sciences Clemson University Clemson, SC, USA 29634-0311.

## Abstract

Probiotic supplementation supports poultry gut health by modulating microbiome and promoting immune development, yet limited information is known about the effects of early, particularly embryonic, supplementation. In this study, we investigated the effects of administering a lactobacilli cocktail *in ovo* (embryonic day 18), post-hatch, or both on gut immunity and the succession of the cecal microbiota in broilers over five weeks. 16S rRNA gene-based sequencing of cecal contents revealed a steady increase in Shannon diversity during the first three weeks (PERMANOVA, *p* < 0.005), with community structure stabilizing by week 3 across all groups. *In ovo* lactobacilli administration improved early hatch rates and modulated microbial composition during early succession, including reductions in *Klebsiella* and *Enterococcus*, and enrichment of *Lactobacillus*, during the first two weeks (MaAsLin2, *q* < 0.25). These microbiome shifts were accompanied by a reduced expression of pro-inflammatory cytokines (IFN-γ, IL-1β, and IL-8) in cecal tonsils. These findings highlight the transient yet critical role of early-life probiotic interventions in shaping gut microbial colonization and immune response in broiler chickens. More importantly, a single *in ovo* lactobacilli dose yielded effects comparable to weekly oral or combined administration.

## Introduction

The avian gut microbiome is a complex and dynamic ecosystem that influences digestion, absorption, immune system development, host metabolism, organ development and overall performance ^1^. In modern broiler production, chicks have no direct contact with adult birds and thus acquire their initial microbiota primarily from the hatching environment. As a result, neonatal chicks are left susceptible to early colonization by opportunistic pathogens such as *Escherichia coli* and *Salmonella* spp., which can compromise gut health during the immediate post-hatch period ^2,3^.

Dietary intervention, including supplementation of probiotics, has been recognized as a strategy to support early gut health and microbiota development ^4–6^. Probiotics are defined as “live, non-pathogenic microorganisms which, when administered in adequate amounts, confer a health benefit on the host.” ^7^ They exert their benefits by competitively excluding pathogens, producing antimicrobial compounds such as bacteriocins, generating organic acids like lactic acid and butyrate, which not only create an unfavorable environment for enteric pathogens but also modulate the host immune system and promote gut homeostasis ^8–10^.

As early colonizers of the poultry gut, lactic acid bacteria (LAB) are recognized for their probiotic properties ^11^. They belong to the phylum Firmicutes, and species such as *L. reuteri*, *L. acidophilus*, *L. crispatus* and *L. animalis* are often isolated from healthy chicken intestines, crop or ceca ^12^. Various LAB strains have been shown to confer protection against enteric pathogens such as *Salmonella*, *E. coli*, *Campylobacter* and *Clostridium perfringens*. For example, in broilers challenged with *Salmonella*, *Lactobacillus rhamnosus* GG significantly reduced pathogen loads in the ceca, liver, and spleen while modulating gut microbiota composition ^13^. *L. acidophilus* has also been shown to enhance gut barrier integrity and reduce inflammatory cytokine expression under *E. coli* challenge ^14^. Similarly, *L. salivarius* significantly reduced cecal colonization by *Campylobacter*, with some strains achieving over 2-log reductions in pathogen load without disrupting microbiome balance ^15^. Several *Lactobacillus* species, including *L. johnsonii*, *L. crispatus*, *L. salivarius*, and *L. reuteri*, have been associated with decreased intestinal lesions, modulation of gut cytokine expression, and stabilization of gut microbiota in response to *Clostridium perfringens* infection ^16^.

As such, these species may serve as suitable probiotics to influence the early succession of the microbiota. While studies have evaluated the early impact of *in ovo* administered LAB on the microbiome of chicks ^17–20^, their influence on weekly microbial succession throughout the rearing period has not been previously established.

Colonization of the chick gut microbiome begins immediately after hatching. Initially, the microbial community is dominated by facultative anaerobes from the environment (such as *Enterobacteriaceae*), but within a week, the succession shifts towards Firmicutes like *Lactobacillus,* with the microbiome stabilizing within the first two to three weeks of age ^21^. A recent study showed that the relative abundance of *Lactobacillus* changed rapidly from day-old chicks (1-6%) to week-old chicks (40%) ^22^. Early colonizers like *Lactobacillus* also engage in molecular crosstalk with the host. The gut microbiome and host immune system communicate through interactions between bacterial components and pattern recognition receptors on the intestinal epithelium and various immune cells ^23–25^.

Manipulation of the gut microbiome of hatched chicks using probiotic lactobacilli has been found to be a useful approach to modulating both gut immunity and microbiome composition ^8^. However, it is widely recognized that continuous administration is typically required to achieve stable and lasting colonization ^26^. This could be attributed to the fact that most of these studies supplement exogenous non-poultry-specific probiotics to the existing microbiome in hatched chicks. As a result, these probiotics may not be able to persistently colonize the intestinal tract and, therefore, exert transient protective effects, requiring continuous supplementation ^27^. Early administration of probiotics through *in ovo* inoculation may enhance the gut microbiome composition of neonatal chicks by seeding the developing gut and competitively excluding pathogens^28–30^.

Given the importance of early microbiome establishment and immune system maturation in newly hatched chicks ^23,30,31^, this study was carried out to evaluate and compare the effects of *in ovo* and post-hatch administration of well-characterized, poultry-derived probiotic lactobacilli on the succession, diversity and taxonomic composition of cecal microbial communities as well as gut immune responses in broiler chicks from hatching to five weeks of age.

## Results

### *In ovo* administration of probiotic lactobacilli positively influenced hatchability outcomes

Poultry-specific probiotic cocktail containing *Lactobacillus reuteri*, *L. crispatus*, *L. animalis*, and *L. acidophilus* (10^7^ CFU/mL) was administered with a dosing volume of 100 µL following ^31^. Group A (control) received no probiotics, Group B received *in ovo* probiotics only, Group C received both *in ovo* and weekly oral probiotic supplementation, and Group D received oral probiotics post-hatch only (Figure 1A). Chicks began hatching on embryonic day 20, with the probiotic-treated group exhibiting a higher early hatch rate (8.89%, 12/135 eggs) compared to the untreated group (4.45%, 6/135 eggs). Overall hatchability was also slightly higher in the probiotic-treated group (99.26%, 134/135 eggs) than in the untreated control group (97.04%, 131/135 eggs), reflecting a 2.2% improvement in hatchability associated with probiotic administration (Figure 1B). To investigate the effect of early *Lactobacillus* supplementation on cecal microbial succession, we performed 16S rRNA gene sequencing on cecal samples collected weekly from four experimental groups of broiler chickens (8 samples/group/week) raised under standard conditions (Figure 1A).

**Figure 1.**
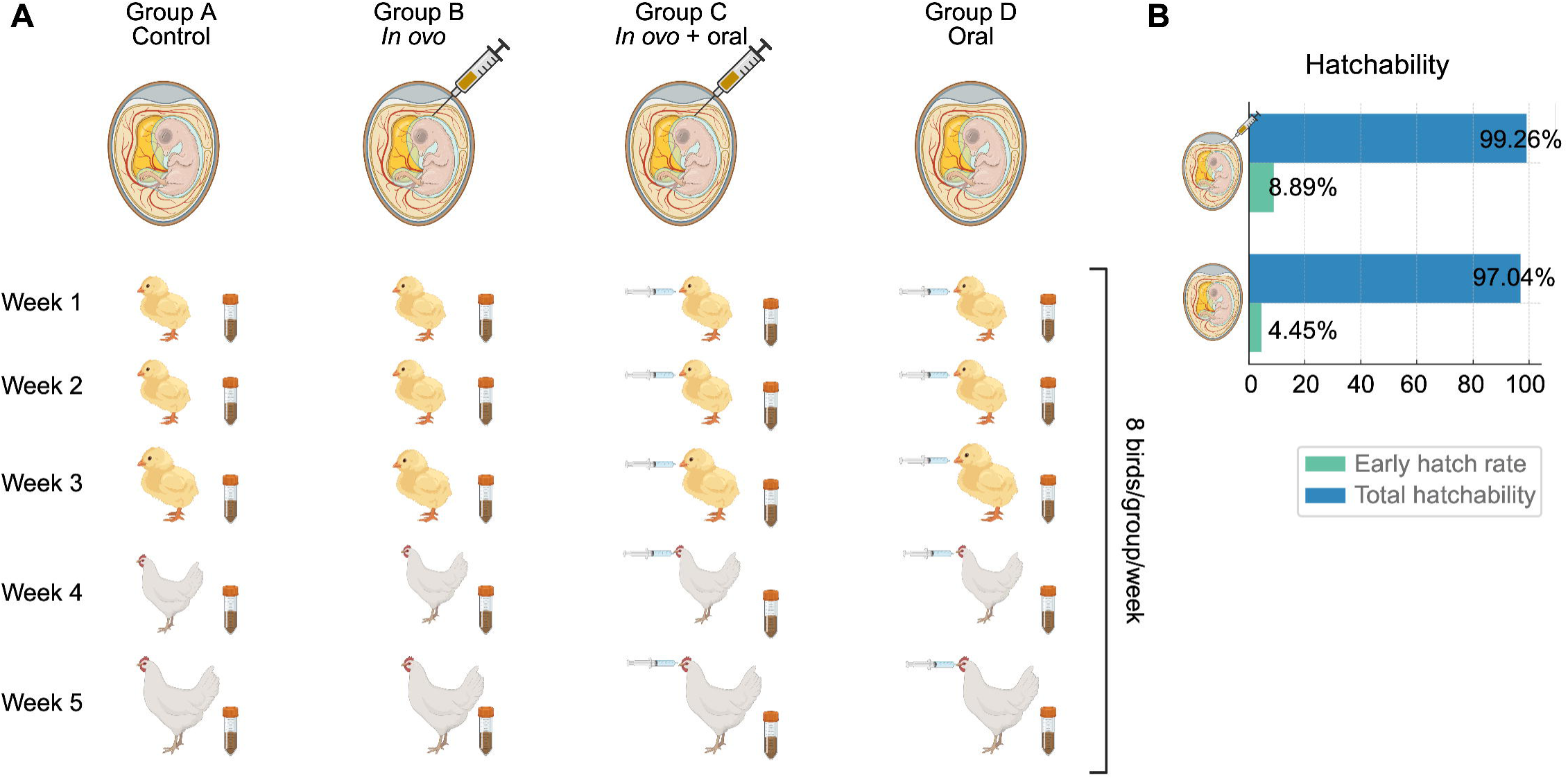
Experimental design of the study and hatchability results.

### Microbial succession in the broiler cecum is marked by increased diversity and Clostridiales dominance

The results revealed that cecal microbiota diversity increased steadily during the first three weeks (Shannon diversity, mean week 1 = 3.00), stabilizing by week three (means in weeks 2, 3, 4, 5 = 3.60, 3.87, 4.07, and 4.00) (Figure 2A). This increase was observed in all four treatment groups, indicating steady microbial succession independent of lactobacilli administration. Similar patterns of microbial structural changes, as assessed by nonparametric multidimensional scaling (NMDS) of the Bray-Curtis distance calculated from ASVs, were also observed over time (Figure 2B). No difference in overall microbiota structure was observed by the end of the experiment (weeks 3 – 5). However, comparison of the microbiota structure at each week demonstrated significant separation of untreated chickens from the remaining treatment groups at weeks 1, 2, and 3, suggesting some effect of lactobacilli manipulation on microbial succession (PERMANOVA, *p* < 0.005).

**Figure 2.**
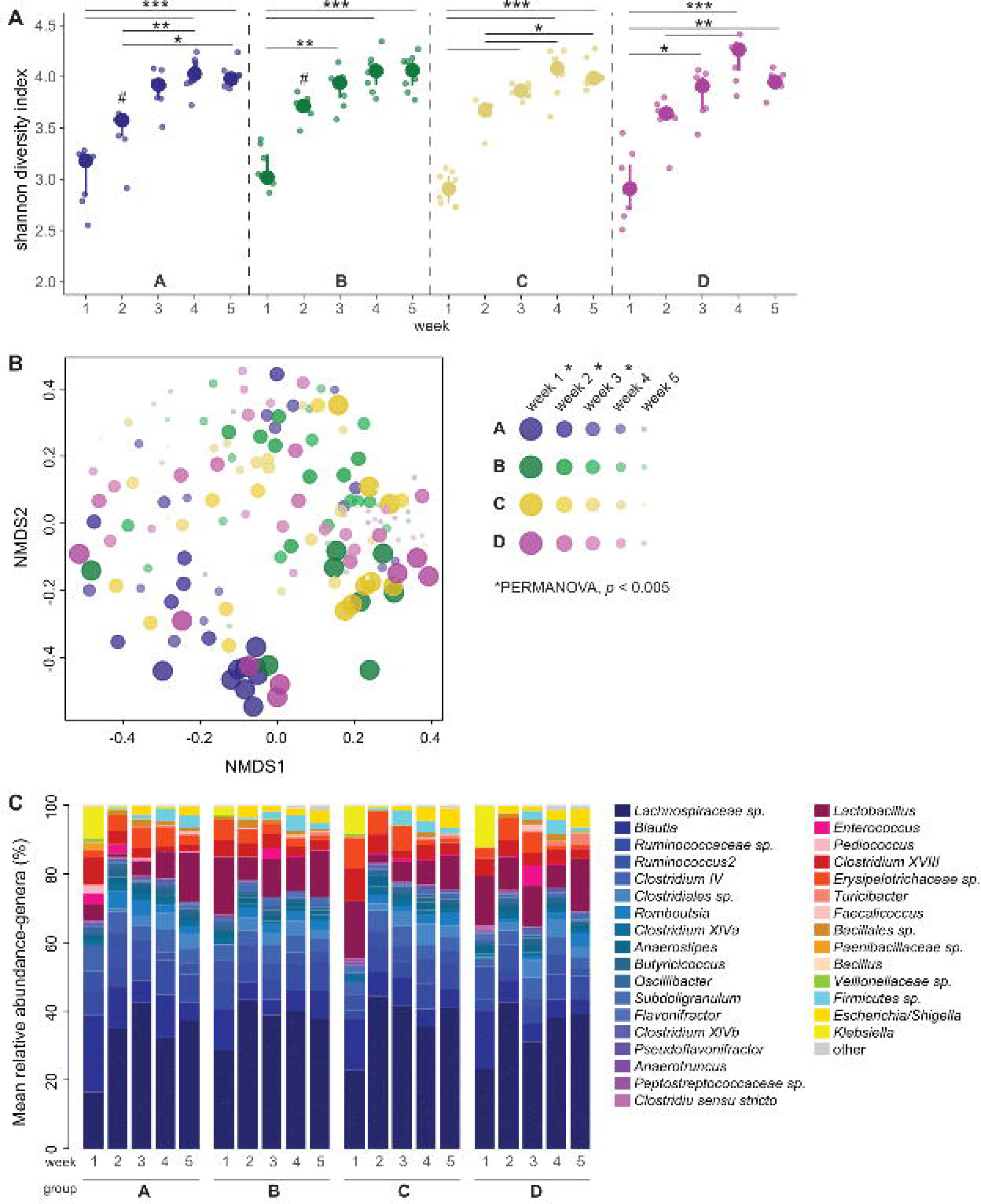
Diversity increases early during microbial succession of the broiler chicken cecum. **A)** Shannon diversity index calculated from amplicon sequence variants (ASVs) in broiler chicken ceca within untreated (group A) chickens or chickens treated with lactobacilli *in ovo* (group B), *in ovo* and orally (group C), or orally (group D). Dunn’s test, \**p* < 0.05, \*\**p* < 0.005, \*\*\**p* < 0.0005 for comparisons within each group; #*p* < 0.05 for comparisons within each week across groups (week 2, A:B groups only significant finding). **B)** Non-metric multidimensional (NMDS) scaling of the Bray-Curtis dissimilarity index calculated from ASVs. PERMANOVA, *p* < 0.001 calculated across groups (A, B, C, and D) within each week. **C)** Mean relative abundance of the most abundant genera (> 2%) in each treatment group, per week.

The overall phyla observed to inhabit the broiler chicken ceca were limited to Bacillota (Firmicutes, overall mean = 96.2%) and Pseudomonadata (Proteobacteria, 3.64%). At the order level, Clostridiales represented the most abundant group (overall mean = 73.64%) across all samples, followed by Lactobacillales (10.5%), Erysipelotrichales (8.85%), and Enterobacteriales (3.64%). There was a slight increase in Clostridiales in all treatment groups from week 1 (mean = 63.9%) to week 2 (80.5%), represented by increases in an unclassified *Lachnospiraceae genus* (week 1 mean = 22.8%, week 2 = 41.3%) and decreases in *Blautia* (week 1 = 16.5%, week 2 = 7.63%) from week 1 to week 2 that stabilized across subsequent weeks. In contrast, Enterobacteriales decreased from week 1 (8.0%) compared to subsequent weeks (means at weeks 2, 3, 4, and 5 = 1.79%, 1.74%, 2.41% and 2.41%). Most of this decrease over time was explained by decreased levels of *Klebsiella* (week 1 = 8.0%, week 5 = 0.008%) and increases in the *Escherichia/Shigella* genus (week 1 = 0%, week 5 = 4.26%).

### Lactobacilli administration influences early microbial succession

Given that our PERMANOVA analysis at weeks 1 and 2 suggested divergent community structures between the treatment groups, we sought to identify compositional factors influencing these differences. We compared genus-level compositional data totaled from individual ASVs to decrease noise at each week. Using MaAsLin2 ^32^, we identified several differentially abundant genera across the weekly comparisons (Figure 3A). Many differentially abundant genera were Clostridiales, the most abundant order identified across the whole dataset; however, these genera did not necessarily demonstrate consistent patterns (i.e., consistent decreases or increases) across subsequent weeks. Several genera displayed consistent differences during weeks 1 and 2. More specifically, *Klebsiella* was significantly decreased in the *in ovo* treatment group (B) compared to the untreated, control group (A), dually treated group (C), or orally treated group (D) (Figure 3B, Dunn’s test, p < 0.05), but decreased rapidly to almost zero in weeks 3, 4, and 5 in all groups. *Enterococcus* was also decreased in all treated groups compared to the control group A in weeks 1 and 2, but increased in all three treated groups in weeks 3, 4, and 5 (Figure 3B, Dunn’s test, p < 0.05). In contrast, *Lactobacillus* increased in all treatment groups compared to the control in week 1 and remained elevated in most of the treatment groups throughout week 4 (Figure 3C, Dunn’s test, p < 0.01). These results suggest that lactobacilli interventions primarily influence lactobacilli and other taxa during early microbial succession.

**Figure 3.**
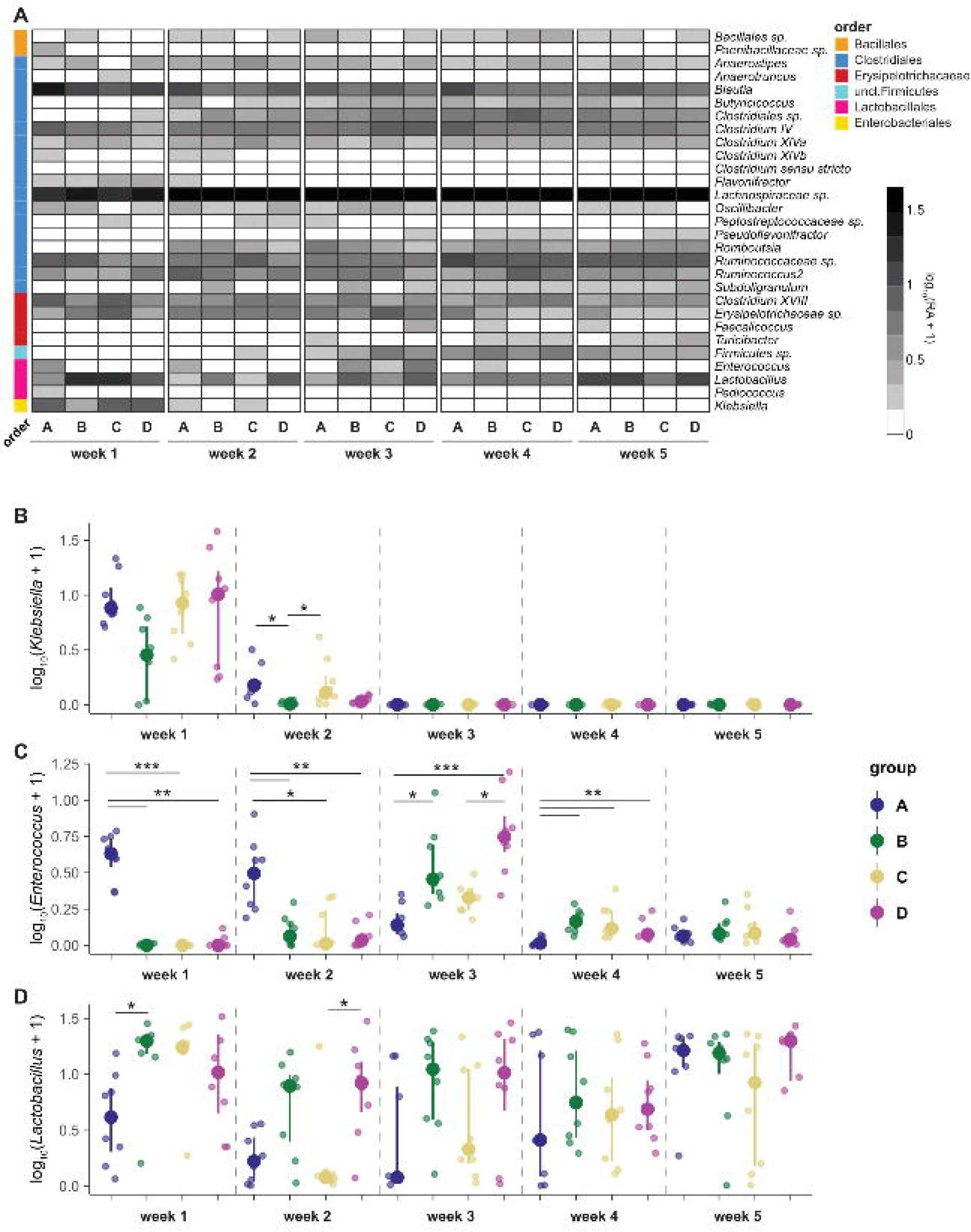
Lactobacilli treatment influences early microbial succession in the broiler chicken cecum. **A)** Relative abundance (log_10_-transformed) of differentially abundant genera across treatment groups (A = control; B = *in ovo*, C = *in ovo* and oral, D = oral lactobacilli) within each week, as identified by MaAsLin2. (Linear model with BH correction, *q* ≤ 0.001). Relative abundance (log10-transformed) of **B)** *Klebsiella*, **C)** *Enterococcus*, and D) *Lactobacillus* in each treatment group by week. Dunn’s test, \**p* < 0.05, \*\**p* < 0.005, \*\*\**p* < 0.005.

### *In ovo* lactobacilli administration alters early gut microbial community structure

To assess the impact of *in ovo* probiotic delivery on early gut community structure, we performed NMDS ordination based on Bray–Curtis dissimilarities. At week 1, NMDS plots revealed clear clustering among treatment groups, and PERMANOVA confirmed a significant treatment effect (F = 1.919, p = 0.001), with treatment explaining 17% of the variance in microbial composition. Furthermore, pairwise PERMANOVA comparisons between the control group (A) and the in ovo group (B) demonstrated a significant shift (F = 1.878, p = 0.001), accounting for 12% of the variation. At week 2, PERMANOVA continued to show significant differences across treatment groups (F = 1.388, p = 0.001), with group-level variance explaining 12.9%. The *in ovo* group (B) remained significantly different from the control (A), with 14.6% of the variance explained (F = 2.394, p = 0.001). Together, these findings indicate that a single *in ovo* dose of probiotics was sufficient to induce early restructuring of the cecal microbiota two weeks post-hatch (Figure 4A).

**Figure 4.**
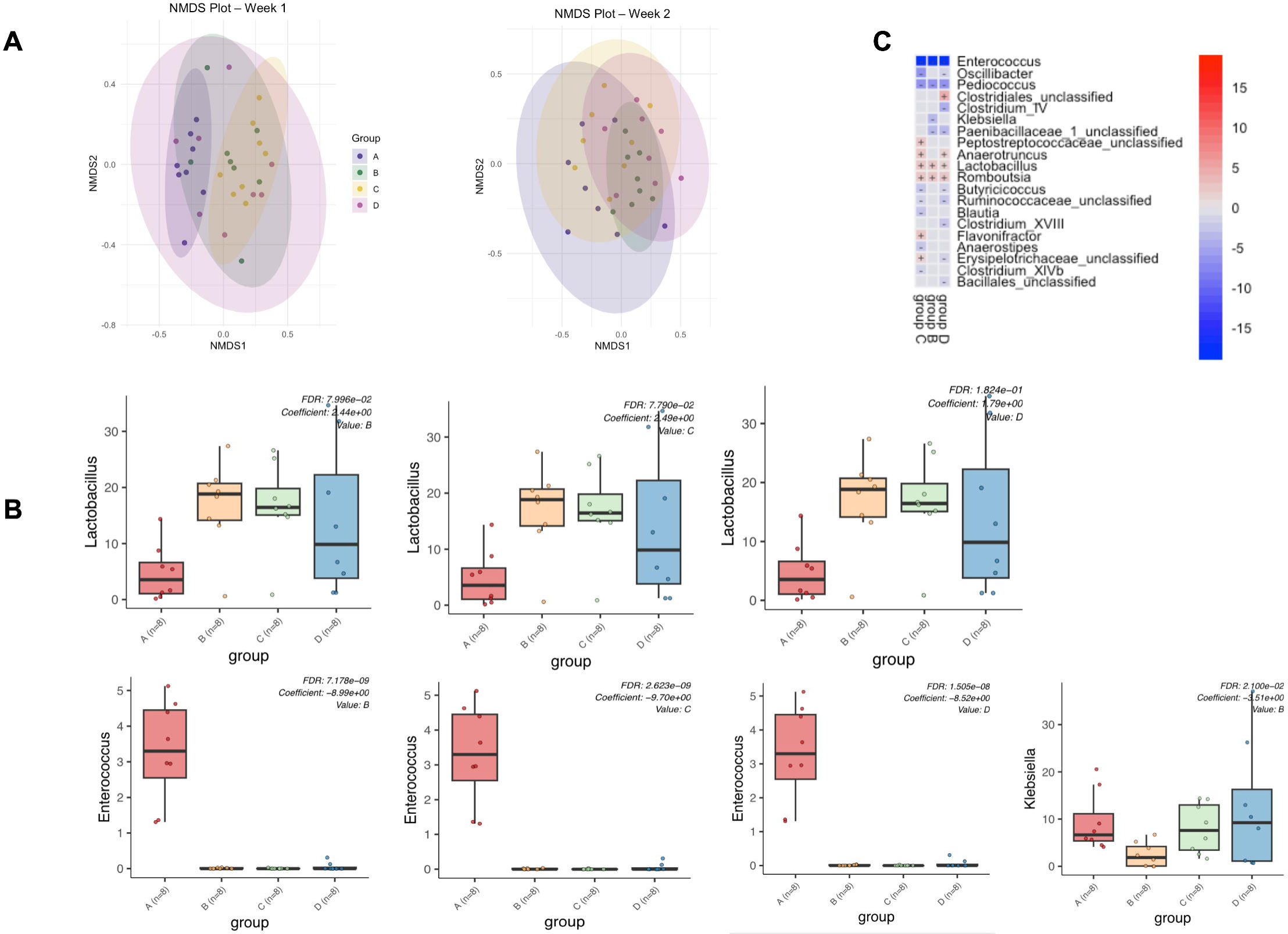
MaAsLin2 analysis reveals early genus-level shifts driven by lactobacilli administration. **A)** Non-metric multidimensional scaling (NMDS) plots of Bray–Curtis dissimilarities showing cecal microbiota structure at week 1 and week 2 across treatment groups. Each point represents an individual sample. **B)** Barplots display genus-level relative abundances of *Klebsiella*, *Lactobacillus*, and *Enterococcus* across treatment groups (A: control, B: in ovo, C: in ovo + oral, D: oral only). Differential abundance testing was performed using MaAsLin2, with group A as the reference. **C)** Heatmap depicting differentially abundant genera across groups B (*in ovo*), C (*in ovo* and oral), and D (oral), compared to the untreated control group A. Genera with positive associations are colored red (increased relative abundance), while blue indicates negative associations (reduced relative abundance), with color intensity representing the signed log-transformed FDR-adjusted p-values (–log(q-value) × sign(coefficient)).

### MaAsLin2 analysis reveals early genus-level shifts driven by lactobacilli administration

To identify microbial taxa associated with lactobacilli administration, we applied MaAsLin2 to genus-level relative abundance data, using the control group (A) as reference. At week 1, MaAsLin2 revealed that *Lactobacillus* was significantly enriched in all probiotic-treated groups, with the highest increase in the *in ovo* group (B) (FDR = 7.996e–02; Coefficient = 1.44; Figure 4B). In contrast, opportunistic genera *Enterococcus* and *Klebsiella* were significantly reduced in all treated groups relative to controls. *Enterococcus* was nearly undetectable in groups B, C, and D but prominent in group A (FDR = 2.623e–09; Coefficient = –9.70), while *Klebsiella* showed the greatest depletion in group B (FDR = 7.996e–02; Coefficient = –2.44). These findings suggest that early-life probiotic administration, particularly via *in ovo*, shows early genus-level shifts.

To further resolve taxonomic drivers of community differences at week 1, we used MaAsLin2 to identify significantly associated genera. The resulting heatmap (Figure 4C) shows differentially abundant genera across the groups B *(in ovo)*, C (*in ovo* + oral), and D (oral), compared to the control (A). *Enterococcus* and *Klebsiella* were significantly reduced, consistent with prior Dunn’s test analysis, while *Lactobacillus*, *Romboutsia*, and *Anaerotruncus* were positively associated with the lactobacilli treatment.

### Lactobacilli administration suppresses early expression of pro-inflammatory cytokines in cecal tonsils

The expression of interferon (IFN)-γ, interleukin (IL)-1β, IL-6, and IL-8 genes in the cecal tonsils (6 cecal tonsils/group/week) was measured from the first week to the fifth week of age using RT-qPCR. During the first week, the expressions of IFN-γ, IL-1β, and IL-8 were downregulated in the oral group (P<0.05, P<0.01, and P<0.01, respectively) compared to the control. (Figure 5) Similarly, in the *in ovo* group, IFN-γ and IL-8 were significantly reduced (P<0.05), while in the *in ovo* and oral combined group, IL-1β and IL-8 were significantly reduced (P<0.05). A lower, but not statistically significant, decrease in the expression of IL-6 was observed in the *in ovo* and oral groups compared to the control group. From weeks 2-5, there were no differences observed in the expression of any of these genes across all treatment groups (data not shown).

**Figure 5.**
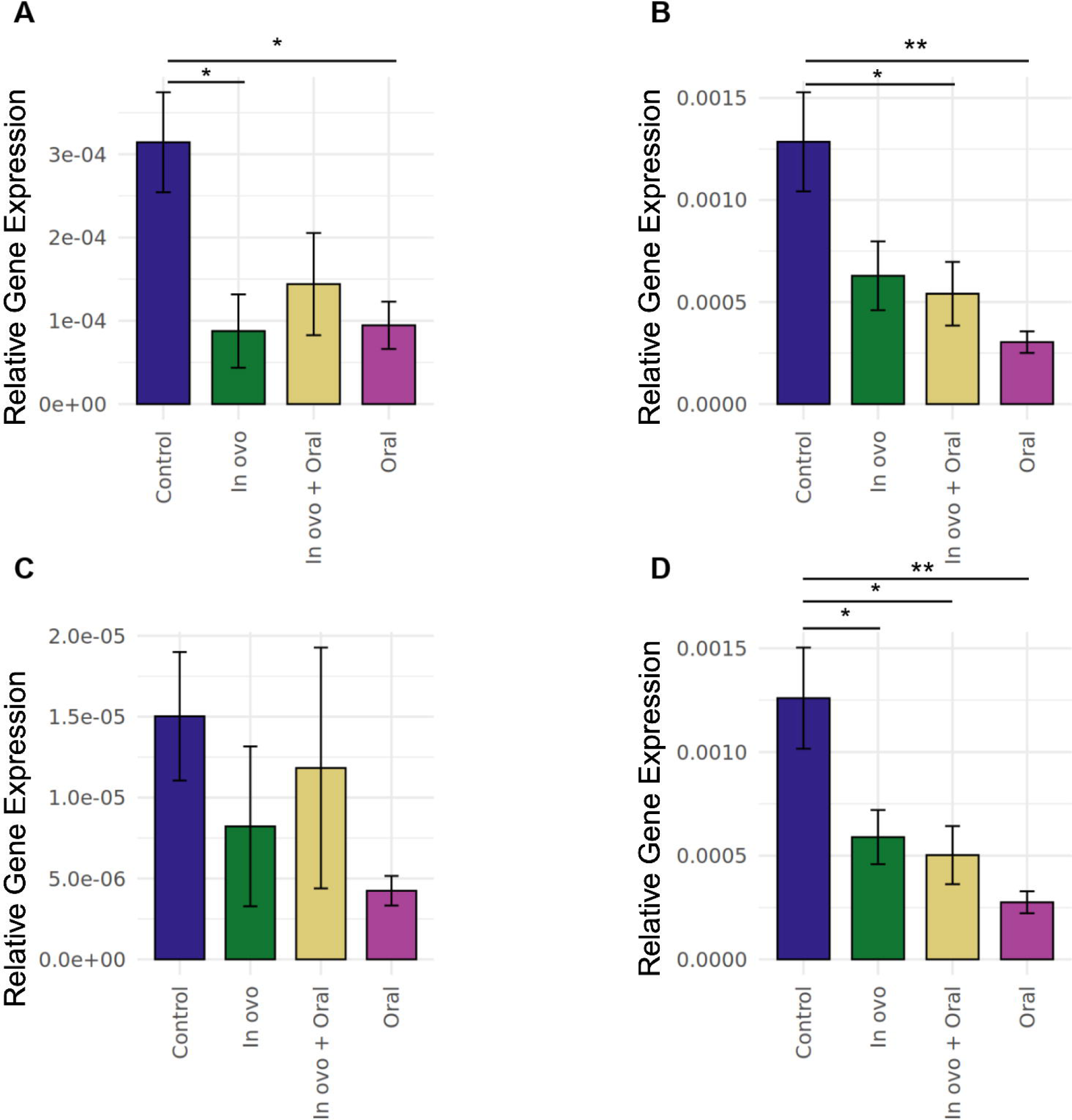
Relative gene expression of proinflammatory cytokines (IFN-γ, IL-1β, IL-6, and IL-8) in the cecal tonsils in the first week of age. The expression of the target genes was calculated relative to the housekeeping gene (β-actin). Statistical significance among treatment groups was calculated using a non-parametric test. Error bars represent the standard error of the mean (SEM). An asterisk (*) indicates a significant difference (P<0.05), a double asterisk (**) indicates a significant difference (P<0.01) and a triple asterisk (***) indicates a significant difference (P<0.001) among the treatment groups.

## Discussion

The chicken gut microbiome is widely recognized to influence gut development, immune function, and colonization resistance against enteric pathogens ^33^. Evidence suggests that early life is important for the acquisition and establishment of the gut microbiome, during which initial microbial colonizers shape gut community structure and the maturation of the gut-associated immune system ^34^. The establishment of a healthy gut microbiome during early life is thus a key determinant of intestinal development, immune maturation, and resistance to pathogen colonization in poultry ^24,33^. Probiotic interventions have been shown to modulate cecal microbial diversity, enhance metabolic activity, and influence mucosal immune responses when administered post-hatch ^35^. However, the extent to which these effects can be enhanced through early manipulation, such as *in ovo* probiotic administration, remains underexplored^2,31,36,37^. In this study, we compare *in ovo*, post-hatch, and combined administration of a lactobacilli cocktail to assess their effects on microbial succession, taxonomic composition, and mucosal immune responses in the chicken cecum.

At the compositional level, our 16S rRNA gene-based study demonstrated that microbial diversity increased steadily over time across all groups, reflecting normal microbial succession in the broiler cecum regardless of the *Lactobacillus* treatment. Birds receiving lactobacilli treatments exhibited earlier and higher colonization by beneficial taxa such as *Lactobacillus*, *Lachnospiraceae*, and *Clostridiales* while showing significantly reduced levels of opportunistic genera like *Klebsiella* and *Enterococcus*. In contrast, the cecal microbiota of untreated birds was initially dominated by *Klebsiella* and *Enterococcus*, which declined over time as microbial diversity increased, an observation consistent with previous reports of early-life microbial succession^23^. Both Enterococcus and *Klebsiella* are frequently isolated from poultry environments such as feed and litter and can form persistent biofilms, exhibit biocide tolerance, and survive harsh conditions ^38–40^. A study found out that multidrug resistant *Klebsiella* had high prevalence up to 35% in retail poultry meat ^41^. Although some *Enterococcus* strains are used as probiotics, many carry antibiotic-resistant genes for tetracycline and beta-lactams ^42^.

Multiple studies have shown that *in ovo* probiotic administration leads to early enrichment of *Lactobacillus* and suppression of opportunistic genera in the gut ^26,27,43,44^. For example, Wilson et al. demonstrated that chicks treated *in ovo* with adult microflora exhibited accelerated establishment of mature microbiota with significantly higher Lactobacillaceae/*Lactobacillus* abundance and decreased abundance of Enterococcaceae/*Enterococcus* at hatch compared to controls ^26^. However, the study monitored microbial dynamics only up to day 10 post-hatch, at which point the observed changes had largely stabilized. In a comparable manner, Arreguin-Naca and colleagues administered a defined adult hen microbiota *in ovo* and observed a significant reduction in Enterococcaceae/*Enterococcus* and Enterobacteriaceae at hatch, accompanied by an increase in beneficial butyrate-producing taxa such as *Ruminococcus* and *Butryricoccus* ^45^.

Likewise, Pedroso et al. used a complex adult-derived competitive exclusion culture and found that probiotic-treated chicks had higher microbial diversity and early colonization by beneficial taxa, aligning with our observations ^46^. Furthermore, studies show that microbiota community differences between treated and untreated birds converge by approximately 3-6 weeks, reinforcing our findings^29,43,47^.

We also observed that *Lactobacillus* itself was a differentiating genus in probiotic-treated vs control birds. Beneficial microbes such as *Lactobacillus* can restrict growth of these opportunistic pathogens through multiple complementary mechanisms in the gut, such as production of inhibitory metabolites, competitive exclusion, enhanced barrier function and modulation of the immune system ^48,49^. Moreover, an increase in *Lactobacillus* is negatively correlated with other genera, limiting their growth likely due to niche competition ^50^. Our findings coincide with previous studies that have also demonstrated the effectiveness of early lactobacilli seeding in promoting rapid gut colonization ^23,46,51,52^.

Additionally, probiotic-treated chicks showed a higher relative abundance of *Clostridiales* compared to controls. Members of this order, including *Lachnospiraceae* and *Ruminococcaceae*, produce butyrate and other short-chain fatty acids that support gut health, nutrient absorption, and barrier integrity ^53,54^. They are typically dominant in mature chicken cecum, suggesting that early probiotic supplementation may accelerate microbial succession toward a more adult-like microbiota.

In addition to the lactobacilli shifts in the microbiome composition, our findings demonstrated a reduction in the expression of proinflammatory cytokines, including IFN-γ, IL-1β, IL-8 and IL-6 in cecal tonsils during the first week of age. These effects may be attributed to the direct action of lactobacilli and/or to lactobacilli-induced alterations in gut microbiome composition, including an increased abundance of butyrate-producing bacteria and the resulting butyrate production, which has been reported to reduce gut inflammation. Aside from their mechanisms of action, prior research has shown that lactobacilli-mediated protection against *Salmonella* is linked to reduced expression of IFN-γ and IL-8 triggered by *Salmonella* infection ^55^. It is therefore plausible to speculate that the decreased expression of these cytokines observed in the cecal tonsils of chicks given *Lactobacillus* either *in ovo* or immediately after hatching may contribute to improved resistance to *Salmonella* during early life. Similarly, a study has shown that dietary supplementation with *Lactobacillus* in broilers has been shown to markedly attenuate *Salmonella*-induced elevations in key inflammatory cytokines, including IL-1β, IL-8, IL-12, and IFN-γ, when compared to untreated, infected controls ^56^. While not investigated in this study, previous research on fecal transplantation in chicks has reported a negative correlation between inflammatory cytokine expression and growth performance ^57^. Therefore, further studies are warranted to evaluate whether *in ovo* lactobacilli supplementation can improve growth performance and enhance resistance to *Salmonella* infection in newly hatched chicks.

Collectively, these results underscore the potential of a single *in ovo* dose of lactobacilli to modulate the intestinal microbiota and the immune system comparable to those achieved through repeated post-hatch dosing. *In ovo* probiotic administration allows for uniform dosing across embryos, facilitating early microbial colonization upon hatch. Previous studies have similarly demonstrated that early microbial seeding can promote mucosal immune development, enhance gut barrier function, and increase resistance to enteric pathogens during the critical first weeks of life ^26,45^ thus reducing the need for continuous supplementation of probiotics post-hatch.

In conclusion, our findings indicate that *in ovo Lactobacillus* supplementation modulated the composition of the cecal microbiota in hatched chicks by enriching the abundance of *Lactobacillus* while reducing the abundance of *Enterococcus* and *Klebsiella*, underscoring the potential of early microbial interventions to shape gut microbial succession. The increased abundance of beneficial taxa alongside reduced inflammatory gene expression further supports the role of early probiotic administration in promoting gut health and immune development. While 16S rRNA gene sequencing offers taxonomic insights, follow-up metagenomic studies are needed to uncover the functional capabilities of early microbial communities.

## Methods

### Ethics statement

All procedures outlined in this study were approved by the Institutional Animal Care and Use Committee (AUP2022-0411) at Clemson University.

### Experimental design

Fertilized broiler eggs (Ross 308 variety) were generously provided by Fieldale Farms, GA, and transported to Morgan Poultry Center (Clemson University, SC). The eggs were incubated in a sanitized egg incubator (GQF Manufacturing Company Inc., GA) with the temperature set to 37°C and humidity maintained between 55-73%, featuring automated turning.

On embryonic day 18 (ED18), the eggs were candled to remove the infertile ones. 110 eggs were randomly selected to receive *in ovo* probiotic administration, and the other 110 were untreated. Sexing of embryos and hatchlings were not performed, and birds used in the study were of mixed sex.

### Preparation of the lactobacilli culture and *in ovo* inoculation

Poultry-specific *Lactobacillus* strains (*L. reuteri*-P43, *L. acidophilus*-P42, *L. animalis*-P38, and *L. crispatus*-C25) were provided by H. M. Hassan’s Laboratory at NC State University. The following preparation of the lactobacilli cocktail was done as per Sharma et al., 2024. A loopful of each frozen *Lactobacillus* species was inoculated into a bacterial culture tube containing 10 mL of MRS (DeMan, Rogosa, and Sharpe) at room temperature. Tubes were incubated overnight at 37°C under anaerobic conditions in a gas jar containing anaerobic packs (BD GasPak™). After 16 hours of incubation, 500 μL of the overnight culture (1%) was inoculated into 50 mL fresh MRS broth and incubated at 37°C under anaerobic conditions for 16-18 hours. The individual tubes were then centrifuged at 4000 rpm for 10 minutes at 4*°*C, and the pellet was washed two times with 1X PBS. After centrifugation, the individual pellets were then resuspended in 1X PBS, and the optical density (OD) of each culture was measured at 600 nm (OD600) using a spectrophotometer (VWR, PA). To determine the number of colony-forming units (CFUs) of each *Lactobacillus* strain, individual growth formulas obtained from the previous study (Sharma et al., 2024) were utilized. All lactobacilli strains were adjusted to a concentration of 10^7^ CFUs/mL in PBS, and a cocktail containing a final concentration of 10^7^ CFUs/mL was used. A final volume of 100 μL was injected.

### Post-hatch procedure

As illustrated in Figure 1, 200 hatched chicks were divided into four groups (n=50): the chicks that received probiotics *in ovo* were randomly divided into two groups: Group B (*in ovo* probiotic supplementation only) and Group C (*in ovo* and oral probiotic supplementation), and the eggs that did not receive probiotics *in ovo* were randomly divided into two groups: Group A (control) and Group D (oral probiotic supplementation only). After hatching, 200 hatched chicks were transported to the Godley-Snell Research Facility (Clemson University, SC) for housing. All birds that received probiotics *in ovo* and post-hatch were housed in separate pens in the same room, while the birds that did not receive any treatments (Group A; control group) were housed in a separate room. Each group of birds was divided into three replicates to minimize the cage effect (n=16-17 per replicate). Chick rearing was done according to the Ross 308 broiler management guidelines by Aviagen®. The birds were fed a standard diet (Purina® Start & Grow® Non-Medicated Chick Feed), and ad libitum water was provided. The birds were raised in floor pens covered with wood shavings. The birds in Groups C and D received oral gavage of the probiotic cocktail containing *L. reuteri, L. crispatus, L. animalis*, and *L. acidophilus* (10^6^ CFUs/mL) on the first day of hatch, and then weekly up to five weeks of age.

### Sample collection

Weekly euthanasia procedures involved randomly selecting and euthanizing ten chickens from each group using CO2 euthanasia over a time period of five weeks. Cecal contents were collected from the ceca in 2 mL tubes and kept in a -80°C freezer (8 samples/group/week). Six cecal tonsils were collected from each group weekly in RNA later and stored in a -20°C freezer for RNA extraction.

### DNA extraction

Total cecal DNA was extracted from 300-400 μL cecal content using the NucleoSpin® DNA soil kit (Macherey-Nagel, Germany). The DNA mass and purity were measured using the NanoDrop One Spectrophotometer (Thermofisher Scientific, MA), ensuring that the 280/260 and 260/230 ratios of the genomic DNA fell within the range of 1.8-2.

### RNA isolation and cDNA preparation

Cecal tonsils were homogenized using the Bead Ruptor Elite (Omni International, GA) for RNA extraction. TRIzol^TM^ (Invitrogen, USA) was employed as per the manufacturer’s instructions to extract total RNA, followed by DNase treatment (DNA-free^TM^ kit, Invitrogen, USA) to get rid of genomic DNA. The Nanodrop One spectrophotometer (Thermo Scientific, USA) was used to evaluate RNA mass and purity. Subsequently, cDNA synthesis was conducted with the Superscript® II First-Strand Synthesis Kit (Invitrogen, USA) and oligo-dT primers (Thermofisher Scientific, USA), following the manufacturer’s protocol. The resulting cDNA was diluted in nuclease-free water (Thermo Scientific, USA) at a 1:10 ratio.

### Quantitative real-time polymerase chain reaction (RT-PCR)

RT-qPCR was performed using the LightCycler® 480 system (Roche Diagnostics). The PCR master mix included 10 µL of SYBR^TM^ Green Master Mix (PowerTrackTM, USA), 1 µL each of forward and reverse primers (10 µM), and 3 µL of nuclease-free water. Each reaction comprised 15 µL of the master mix and 5 µL of cDNA in a 96-well PCR plate (USA Scientific, USA).

The PCR cycling protocol involved an initial denaturation step at 95°C for 5 minutes, followed by 45 amplification cycles. Each cycle consisted of 10 seconds at 95°C for denaturation, annealing (according to the target primers specified in Table 1), and extension at 72°C for 10 seconds. All primers used in this study were synthesized by Sigma-Aldrich (St. Louis, MO). The mRNA expression levels of the target genes were normalized to the housekeeping gene (β-actin) using the Roche LightCycler 480 software, based on the 2^-ΔΔCT^ method (Livak & Schmittgen, 2001).

**Table 1.**
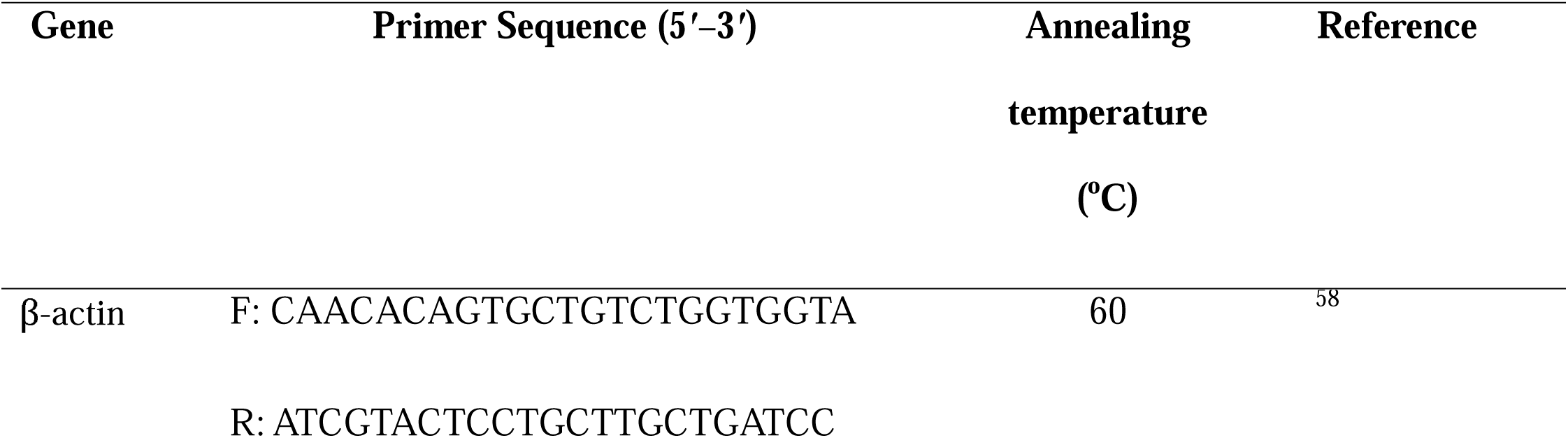
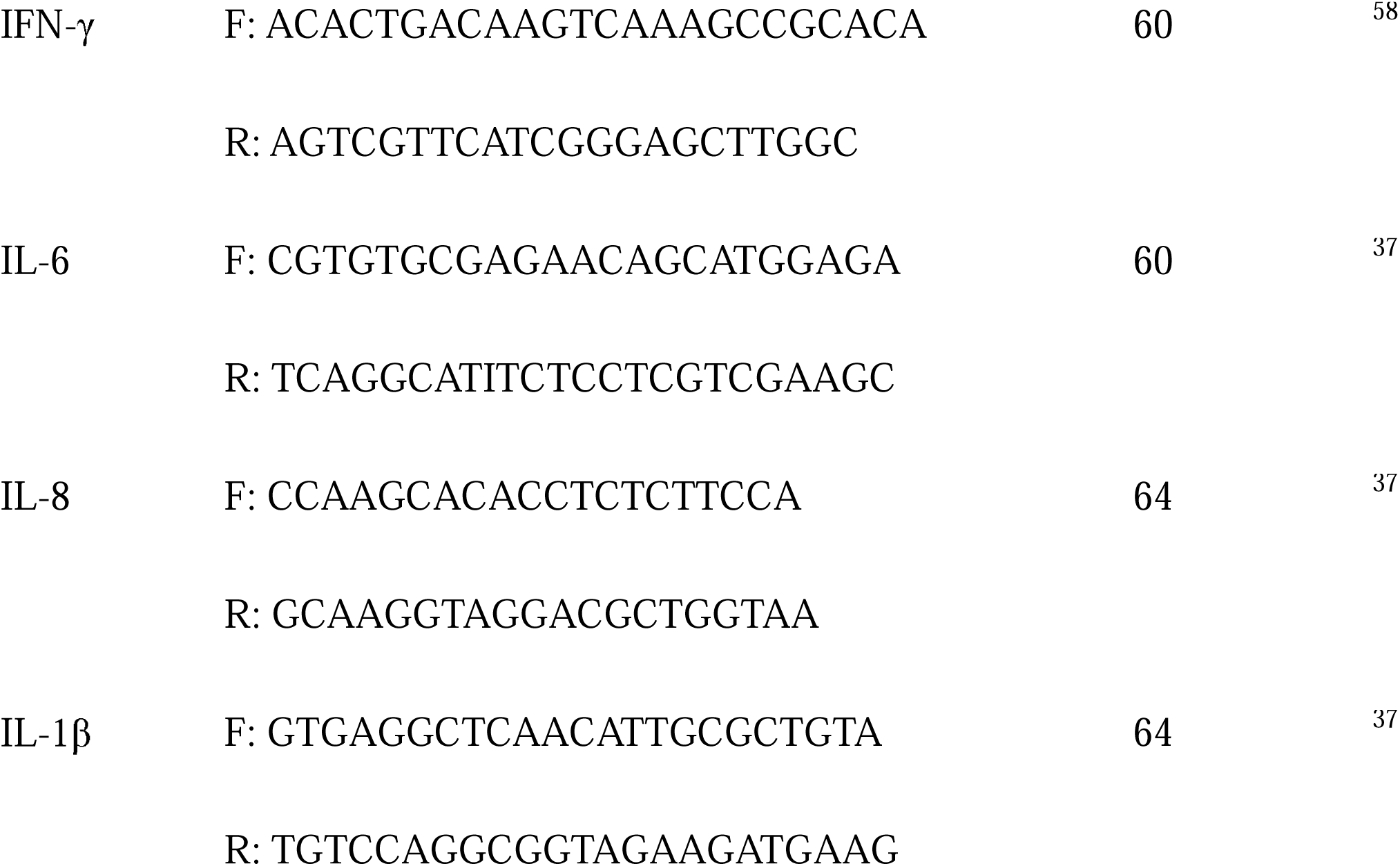
Primer sequences used for real-time quantitative PCR.

### 16S rRNA library preparation

Genomic DNA was diluted to a concentration of 10 ng/µL per well in a volume of 10 µL in a 96-well PCR plate and submitted to the Genomic Sciences Laboratory at North Carolina State University in Raleigh, NC, for library preparation. The V3-V4 hypervariable region of the 16S rRNA gene of the bacterial DNA was amplified using the following forward primer pairs: 16SF: 5’ -TCG TCG GCA GCG TCA GAT GTG TAT AAG AGA CAG CCT G GGN GGC WGC AG - 3’ and 16SR: 5’ - GTC TCG TGG GCT CGG AGA TGT GTA TAA GAG ACA GGA CTA CHV GGG TAT CTA ATC C - 3’. The PCR cycling conditions included an initial denaturation step at 95°C for 3 minutes, followed by 25 cycles of denaturation at 95°C for 30 seconds, annealing at 55°C for 30 seconds, and extension at 72°C for 30 seconds. A final extension step was performed at 72°C for 5 minutes. Following the amplification cycles, the reaction mixture was held at 4°C until further processing. After PCR amplification, the PCR products were purified using AMPure XP bead cleanup (Beckman Coulter Life Sciences). The DNA samples were quantified and normalized to a loading library concentration of 4 nM. The amplicons were subjected to paired-end sequencing on an Illumina MiSeq platform using the MiSeq V3 300x2 sequencing kit. The sequencing was performed with ≥ 25% PhiX control spike in. Paired-end reads of all FASTQ files are available in the Sequence Read Archive (SRA) under BioProject PRJNA1205772.

### Bioinformatic analysis

A total of 26,212,889 raw sequencing reads were generated in fastq.gz format. Demultiplexing was performed using bcl2fastq2 Conversion Software v2.20 (Illumina). Raw sequences were processed using mothur v.1.45.2, following standard operating protocol (Kozich et al., 2013). Paired-end reads were merged into contigs, quality-filtered (maxambig = 0, maxlength = 475, maxhomop = 8), and aligned to the SILVA reference database v132 trimmed to the V3–V4 region. Sequences were then dereplicated and pre-clustered. Chimeric sequences were identified and removed using VSEARCH algorithm. Taxonomic classification was performed using the RDP classifier with the RDP training set v16. Lineages associated with mitochondria, chloroplasts, unknown, Archaea, and Eukaryota were removed. A total of 459,228 Amplicon Sequence Variants (ASVs) were generated, of which low-abundance ASVs (<5 reads in the total dataset) were filtered out, retaining 16,489 ASVs for downstream analysis.

After preprocessing the data in mothur, phyloseq (v.1.48.0), ggplot2 (v. 3.5.1), dplyr (v.1.1.4) and vegan (v.2.6) packages in R software (v.4.4.0) were used. Alpha diversity was assessed using the Shannon index. Kruskal-Wallis test was used for initial comparison among groups, using a Dunn’s post hoc test. Beta diversity was assessed using the Bray-Curtis dissimilarity for differences in microbial community composition across groups from the vegan R package (Oksanen et al., 2024). PERMANOVA was conducted using the adonis function from the vegan package in R. Multivariable Association with Linear Models (MaAsLin2) was used to calculate differentially abundant ASVs across treatment groups within each week (Mallick et al., 2021). Heatmaps of the relative abundances of dominant ASVs at the family level were created using the pheatmap R package, using a log_10_ transformation before plotting relative abundance.

### Gene expression analysis

For the gene expression study, data were analyzed, and graphs were created using R software (v.4.4.0). A non-parametric test (Kruskal-Wallis test) was used, and the results were considered significant if P<0.05. Data are shown graphically as the mean of the relative gene expression data (2^−ΔΔCt^) ± the standard error of the mean (SEM).

## Data availability statement

The 16S rRNA sequencing data generated during this study have been deposited in the NCBI Sequence Read Archive (SRA) under BioProject accession number PRJNA1205772.

## Competing interests

The authors declare no competing interests.

## Funding Declaration

This research was supported by the USDA National Institute of Food and Agriculture Hatch Project SC-1700628 (Accession Number 7004405, Technical Contribution No. 7456), South Carolina Department of Agriculture (ACRE CGP) and Clemson University’s R-Initiatives for funding.

## Acknowledgments

The authors also acknowledge Fieldale Farms Corporation for providing fertilized Ross 308 broiler eggs and the staff at Godley Snell Research Facility and Morgan Poultry Center for assistance with bird rearing. Data analysis was conducted using Clemson University’s Palmetto HPC System, which was made possible with support from the Clemson University Genomics and Bioinformatics Facility (CUGBF), which receives support from the College of Science and two Institutional Development Awards (IDeA) from the National Institute of General Medical Sciences of the National Institutes of Health under grant numbers P20GM146584 and P20GM139769.

## Notes

### Competing Interest Statement

The authors have declared no competing interest.

